# Structure-based engineering of Tor complexes uncovers different roles of two types of yeast TORC1s

**DOI:** 10.1101/2023.07.04.547620

**Authors:** Yoshiaki Kamada, Chiharu Umeda, Yukio Mukai, Hokuto Ohtsuka, Yoko Otsubo, Akira Yamashita, Takahiro Kosugi

## Abstract

Certain proteins assemble into diverse complex states, each having a distinctive and unique function in the cell. The target of rapamycin complex 1 (TORC1) plays a central role in signaling pathways for cells to respond to their environment, such as nutritional status. TORC1 is widely recognised for its association with various diseases. The budding yeast *Saccharomyces cerevisiae* has two types of TORC1s comprising different constituent proteins, Tor1- and Tor2-containing TORC1s but are considered to have the same function. Here, we rationally redesigned the complex states by structure-based engineering and constructed a Tor2 mutant to form TORC2 but not TORC1. Functional analysis of the mutant revealed that the two types of TORC1s induced different phenotypes-rapamycin, caffeine and pH dependences of cell growth and replicative and chronological lifespans. These findings are expected to provide further insights into various fields such as molecular evolution and lifespan.

## Introduction

Various proteins assemble into complex states within cells, and a substantial proportion alters combinations of constituent proteins to proficiently exert their functions in the appropriate spatiotemporal context. Target of rapamycin (Tor), an evolutionarily conserved protein kinase, plays a pivotal role in eukaryotic cell signalling pathways. It responds to changes in the extracellular environment, such as changes in nutritional status, and is associated with various diseases and lifespans (Liu & Sabatini, 2020; Loewith & Hall, 2011). Tor forms two distinct complex dimers, Tor complex 1 and 2 (TORC1 and TORC2) (Loewith *et al*, 2002). They comprise different several constituent proteins, resulting in exerting different functions (Fig. 1A) (Liu & Sabatini, 2020; Loewith & Hall, 2011). In *S. cerevisiae*, a serine/threonine protein kinase, Tor1, assembles into TORC1 with the main partner Kog1 (yeast counterpart of Raptor) and other partner proteins such as Lst8. Another kinase protein, Tor2, assembles into not only TORC1 but also TORC2 with the main partner Avo3 (yeast counterpart of Rictor) and other proteins, e. g. Lst8, Avo1 and Avo2 (Loewith *et al*., 2002). Namely, there are two types of *S. cerevisiae* TORC1s which are composed of Tor1 or Tor2 (hereafter referred to as Tor1-TORC1 and Tor2-TORC1), whereas other species, such as mammals and the fission yeast, *Schizosaccharomyces pombe*, have only one type of TORC1 (Fig. 1A) (Otsubo *et al*, 2017).

**Fig. 1.**
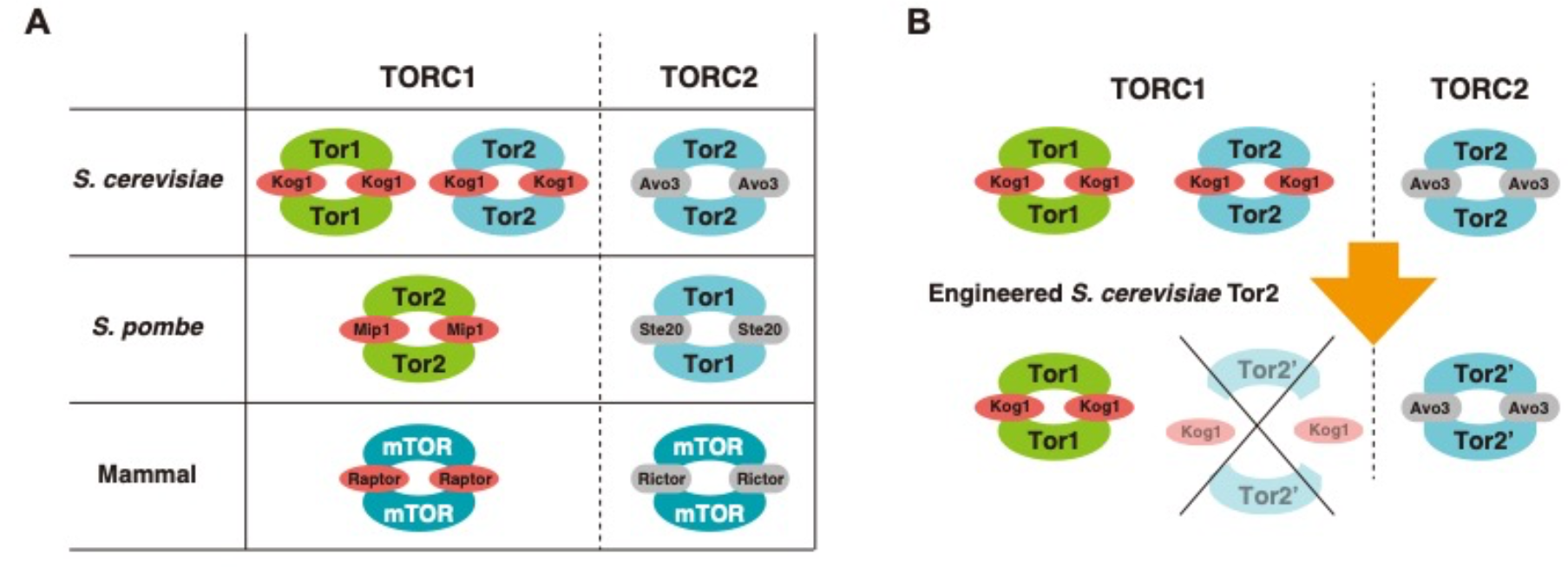
Strategy to compare two types of TORC1 in *S. cerevisiae*. **A.** Orthologue proteins which constitute TORC1s and TORC2s in several species. Only *S. cerevisiae* has two types of TORC1, namely Tor1- and Tor2-containing TORC1s (Tor1-TORC1 and Tor2-TORC1). **B.** Tor2-TORC1 in *S. cerevisiae* is deleted by engineering Tor2 to maintain the binding ability with Avo3 and not with Kog1. In these figures, Lst8 is omitted.

To the best of our knowledge, functional differences between Tor1-TORC1 and Tor2-TORC1 have not yet been studied. Tor2-TORC1 has been ignored because the amount of Tor2-TORC1 in cells is lower than that of Tor1-TORC1 (Loewith *et al*., 2002; Reinke *et al*, 2004). Moreover, the functions are thought to be the same because Tor1 is a homologue of Tor2 with high sequence identity (66.2%) and the same ligands are phosphorylated by both Tor1-TORC1 and Tor2-TORC1 (Kunz *et al*, 1993). However, these complexes exhibit several interesting assembly properties. Tor2 assembles into both complex states, whereas Tor1 does not assemble into TORC2. There is also no chimeric TORC1 dimer which contains both Tor1 and Tor2; in a TORC1 dimer, only either Tor1 or Tor2 is included (Loewith *et al*., 2002; Reinke *et al*., 2004; Takahara *et al*, 2006).

TORC1 and TORC2 activity in *S. cerevisiae* is essential for cell growth (Loewith & Hall, 2011). *TOR1* deletion is not lethal because only Tor1-TORC1 is deleted and Tor2-TORC1 remains. The *tor1*Δ strain has been under intense investigation. It has been shown that the cell lifespan of *tor1Δ* cells is extended (Kaeberlein *et al*, 2005), because partial inhibition of TORC1 mimics calorie restriction, an important factor of longevity (Lin *et al*, 2000). Inhibition of TORC1 using drugs, such as rapamycin or Torin-1, also leads to longevity in various organisms, including yeast, nematodes, flies, and rodents. (Dabrowska *et al*, 2022; Folch *et al*, 2018; Harrison *et al*, 2009; Martinez-Miguel *et al*, 2021; Ohtsuka *et al*, 2021b; Rodríguez-López *et al*, 2020). For lifespan, Tor1-TORC1 is expected to have a function similar to that of TORC1s in other species. However, the function of Tor2-TORC1 function is unclear. *TOR2* deletion is lethal because only Tor2 assembles into TORC2 (Kunz *et al*., 1993; Loewith *et al*., 2002); this could be one of the reasons why Tor2-TORC1 has never been studied.

Studies on Tor complexes by domain exchange did not show the differences between Tor1-TORC1 and Tor2-TORC1; they performed the experiments to identify important interactions for assembling complex states (Hill *et al*, 2018; Tsverov *et al*, 2022). If high-resolution three-dimensional structures are solved using X-ray or cryo-EM, important interactions can be uncovered. Moreover, a comparison of the three-dimensional structures of *S. cerevisiae* Tor1-TORC1 and Tor2-TORC1 may provide clues about their functions. However, high-resolution structures have not yet been obtained.

Recently, remarkable development of computational protein structure prediction and protein design methods has been achieved (Baek *et al*, 2021; Dauparas *et al*, 2022; Huang *et al*, 2016; Jumper *et al*, 2021). Using computational methods, native proteins have been redesigned and their functions have been successfully controlled. For example, artificial activation or inactivation of G protein-coupled receptors and cyclic GMP-AMP synthase by state-targeting stabilisation have been reported (Chen *et al*, 2020; Dowling *et al*, 2023). We have also previously controlled the concerted function, rotation, of a rotary molecular motor, V_1_-ATPase, using a novel approach based on computational protein design methods (Kosugi *et al*, 2023). Here, based on predicted structural models for protein complexes whose experimental structures are unavailable, we engineered a constituent protein to change the pattern of possible combinations and attempted to uncover the biological functions of a protein complex in cells.

In this study, we designed a Tor2 mutant protein which could not form TORC1 but could form TORC2 (Fig. 1B) by structure-based engineering. Mutant strains of *TOR2* showed differences for several phenotypes with those of the *tor1*Δ strain-rapamycin, caffeine and pH dependences of cell growth and replicative and chronological lifespans. These results revealed that the several characteristics of Tor2-TORC1 were different from those of Tor1-TORC1. Based on the differences in their roles, we propose new perspectives for research on the molecular evolution and lifespan.

## Results

### Structure-based engineering of Tor2 to lose its ability to assemble to TORC1, but retain its ability to maintain the TORC2 assembly

To engineer Tor2 not to assemble to TORC1 but to maintain the TORC2 assembly based on the structures, we focused on the interactions between Tor2 and the unique components of each complex–Kog1 for Tor2-TORC1 and Avo3 for TORC2. We aimed to eliminate the interaction of Tor2 with Kog1 and retain it with Avo3 by structure-based engineering. Therefore, reasonable structures for Tor2-TORC1 and (Tor2-)TORC2 are required. However, high-resolution structures of Tor2-TORC1 and TORC2 from *S. cerevisiae* have not been reported, except for a recently resolved cryo-EM structure of the TORC1 inactive condensate, TOROID (Prouteau *et al*, 2023). Both cryo-EM structures of mTORC1 and mTORC2 have been reported at approximately 3.2 Å resolution (Scaiola *et al*, 2020; Yang *et al*, 2017). Therefore, by superimposing homology models of each constituent protein (Tor2, Kog1, and Avo3) from *S. cerevisiae* on the human Tor complex structures, we computationally modelled a dimer of the Tor2 and Kog1 protein complex as Tor2-TORC1, and a dimer of the Tor2 and Avo3 protein complex as TORC2 (Fig. 2). By comparing these two model complex structures, we found a design target region in Tor2 that interacted with Kog1 but not with Avo3; note that other constituent proteins, LST8 and Avo2, also do not interact with this region. This region contacts loop structures of Kog1 which are not in orthologues from other species, *S. pombe* or humans (the sequence alignments are shown in Appendix Fig. S1); this loop structures are expected to contribute to TORC1 assembly of *S. cerevisiae* Tor2. In the design target region of Tor2, nine amino acid residues (A740, A742, K768, A772, A775, A777, L781, F817, and K818) within the HEAT domain were selected and mutated to crash with the characteristic loop region of Kog1 and to stabilize the surface exposed to solvent; hydrophobic and charged residues were mutated to a larger and hydrophilic residue, glutamine, and to larger amino acids with the same charge, respectively. As shown in Fig. 3A, seven combinations of the mutations—K1, K2, K3, K12, K13, K23 and K123—were experimentally validated.

**Fig. 2.**
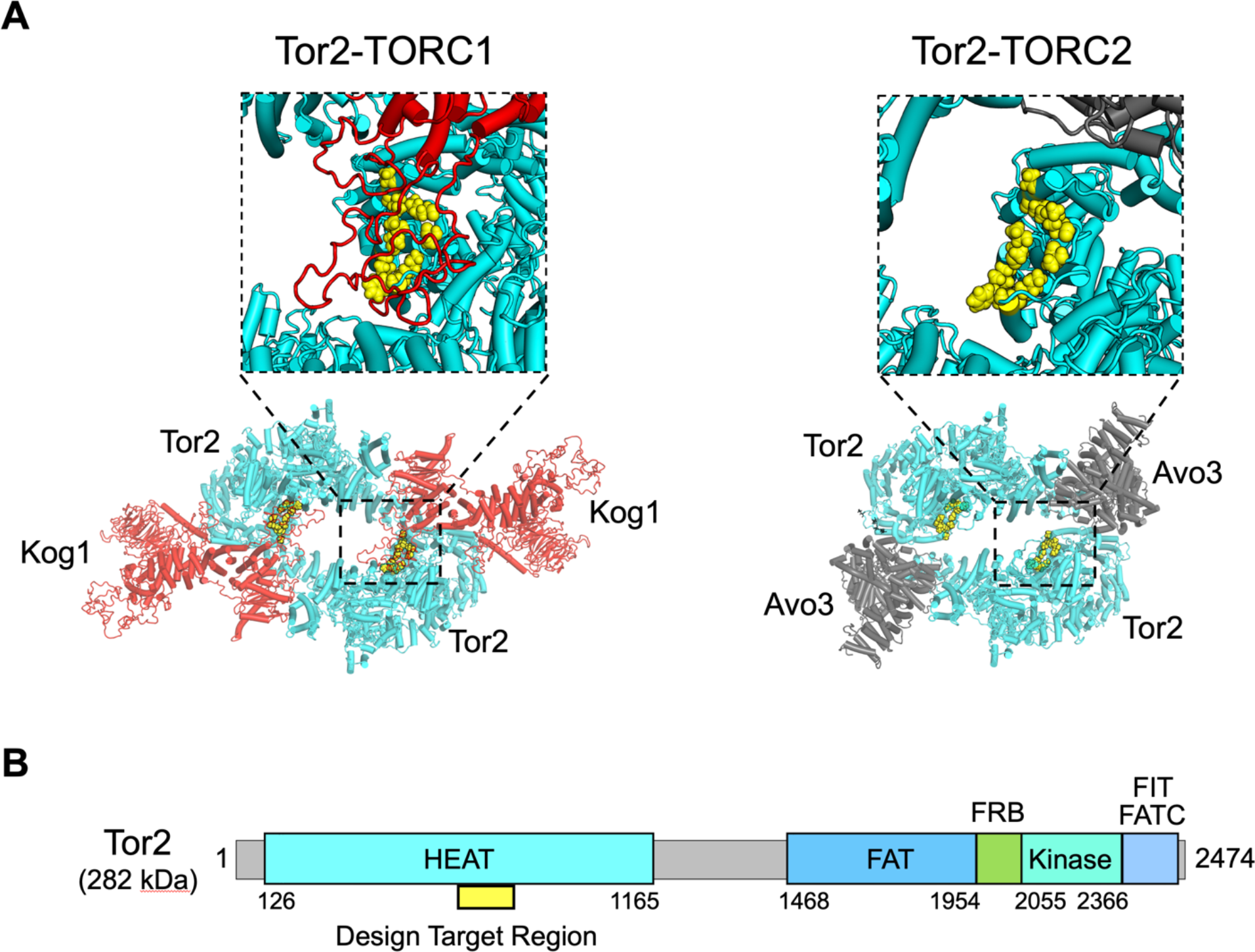
Design target region in Tor2 found by comparing two model structures. **A.** Model structure of Tor2-TORC1 (left) and Tor2-TORC2 (right) (Lst8 is omitted). The amino acid residues in the design target region, with which Kog1 interacts but Avo3 does not, are shown as yellow spheres. **B.** Design target region (yellow) on the primary sequence of Tor2. The region is on the Tor2 HEAT domain.

**Fig. 3.**
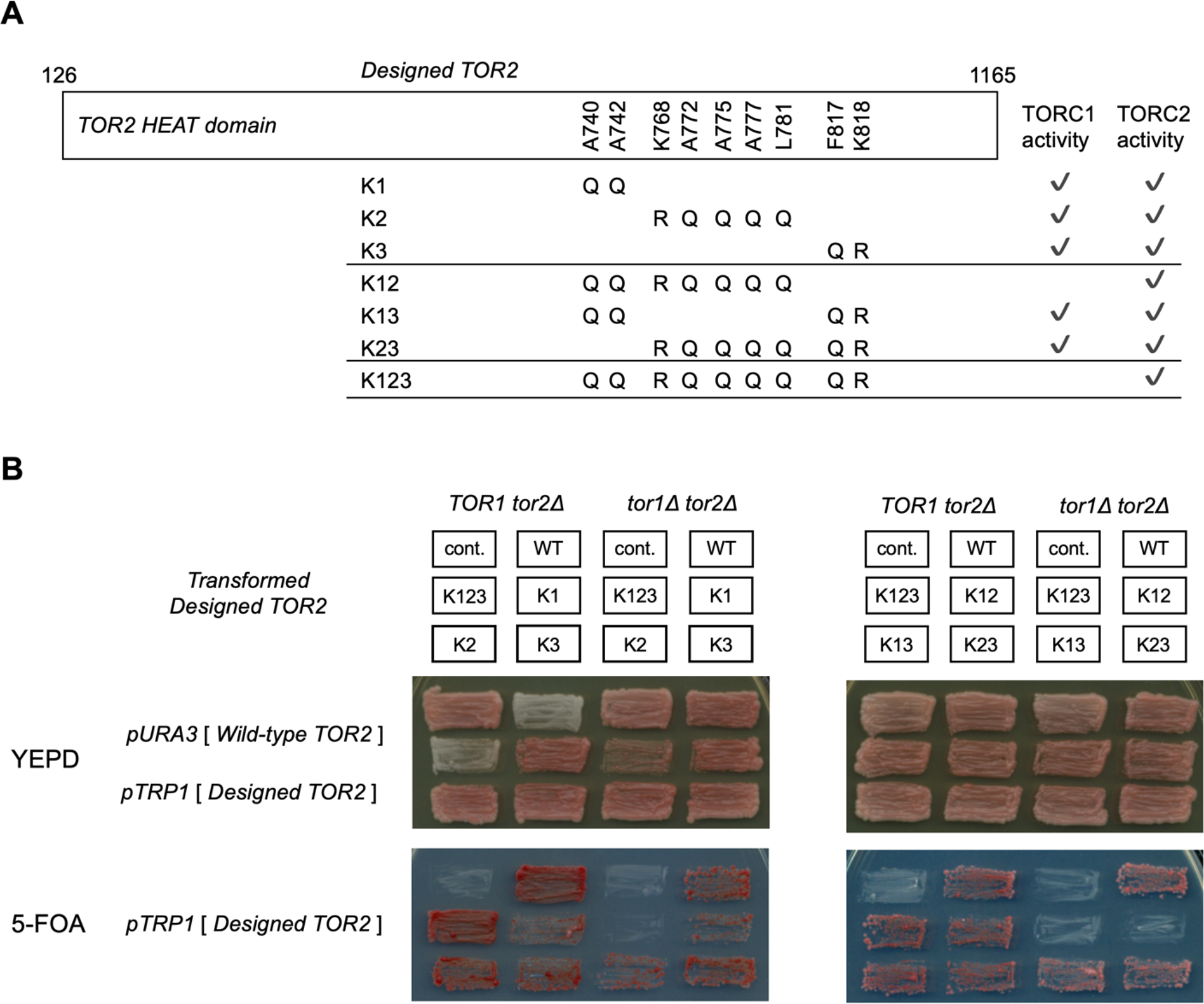
Cell-based assay for the designed Tor complexes having no TORC1 function and retaining TORC2 function. **A.** Mutation site and amino acid type of designed Tor2s. A summary of the cell-based assay is also shown. **B.** Cell-based activity assay of Tor complexes with several designed *TOR2* mutants; K1, K2, K3, K12, K13, K23, and K123. Cells were patched onto YEPD (top) and 5-FOA (bottom) plates and incubated for 2 days at 30 °C. K12 and K123 have no TORC1 activity and retain TORC2 activity, respectively, as designed.

### Engineered Tor2 mutant maintains TORC2 activity but does not have TORC1 activity

Tor2-TORC1 and TORC2 activities in the seven mutant strains were verified using a cell-based assay. The TOR2 mutant plasmids cloned into pRS314 (TRP1) vector were transformed into *TOR1 tor2*Δ and *tor1*Δ *tor2*Δ strains, harbouring the pRS316(URA3)[TOR2] plasmid. The transformants were streaked onto YEPD (control) or 5-FOA plates to select a *ura^-^* cell which loses the URA3-maker wild-type TOR2 plasmid and harbuored only the mutated TOR2 plasmid. Growth on 5-FOA plates was used to evaluate the function of TORC2 (*TOR1 tor2*Δ strain) and TORC1 TORC2 (*tor1*Δ *tor2*Δ strain) (Fig. 3B) because the loss of either TORC1 or TORC2 activity is lethal for cells. For example, K1, K2, K3, K13, and K23 transformants grew on 5-FOA plates in both *TOR1 tor2*Δ and *tor1*Δ *tor2*Δ background as well as the wild type (Fig. 3A and B). In contrast, the K12 and K123 transformants grew on 5-FOA only in *TOR1 tor2*Δ background, indicating that these two *TOR2* mutants do not function as TORC1. These results suggest that the two strains, K12 and K123, exhibit activities as expected from the design.

To further characterise the Tor2(K12) and (K123) mutants, their TORC1 and TORC2 complex-forming abilities were evaluated by co-immunoprecipitation (Fig, 4A-C). Analysis of Tor2-TORC1, in which the HA-tagged Tor2(K12) mutant was pulled down together with FLAG-tagged Kog1, indicated that the Tor2(K12) mutant largely loses its ability to form a Tor2-TORC1 complex with Kog1. However, analysis of TORC2 by FLAG-tagged Avo3 together with cell-based assays indicated that the K12 mutant maintained sufficient Tor2-TORC2 forming ability to function as TORC2. When the same amount of TORC2 was immunoprecipitated for the *in vitro* TORC2 kinase assay, Tor2-TORC2 kinase activity was found to be similar between the wild-type and Tor2(K12) mutant, confirming that mutation sites in K12 did not affect the specific activity of TORC2 (a similar result was obtained by another method using ATPγS as a substrate, as shown in Appendix Fig. S2). These results show that the Tor2(K12) mutant was successfully designed as we expected; in Tor2(K12) mutant cell, Tor2-TORC1 formation is largely compromised while (Tor2-)TORC2 formation is well conserved. HA-tagged Tor2(K123) protein was barely detected in the cell lysate, while it was detected in denatured conditions (Appendix Fig. S3); therefore, we could not perform co-immunoprecipitation analysis. This complex is probably more fragile than the Tor2(K12) mutant complex, although it form the TORC2 and has TORC2 activity in cells. Therefore, for further experiments, we focused on the Tor2(K12) mutant and investigated functions of the two types of TORC1s by comparing the K12 strain (Tor2-TORC1 was almost lost in the cell) with the *tor1*Δ strain (Tor1-TORC1 was lost).

### Phenotypes of *tor1*Δ and *tor2* mutant strains are different from each other

First, the *in vivo* TORC1 kinase activities in *tor1*Δ and *tor2*(K12) strains were estimated by the phosphorylation states of TORC1 substrates, Atg13 or Sch9 (Kamada *et al*, 2010; Urban *et al*, 2007). In both mutant strains, the activity of TORC1 was similar to that of wild-type strain (Fig. 4D). Therefore, even if either of TORC1s is lost, the TORC1 kinase activity of the major ligands is maintained. Incidentally, the TORC2 activity, kinase activity for Mpk1 (Fig. 4E) in cells, and actin organisation in cells (Kamada *et al*, 2005) (Fig. 4F) were also completely maintained. Therefore, Tor2(K12)-TORC2 did not affect the phenotypes observed at 30 °C. The *tor2*(K12) strain showed growth defects at 37°C (Appendix Fig. S4).

**Fig. 4.**
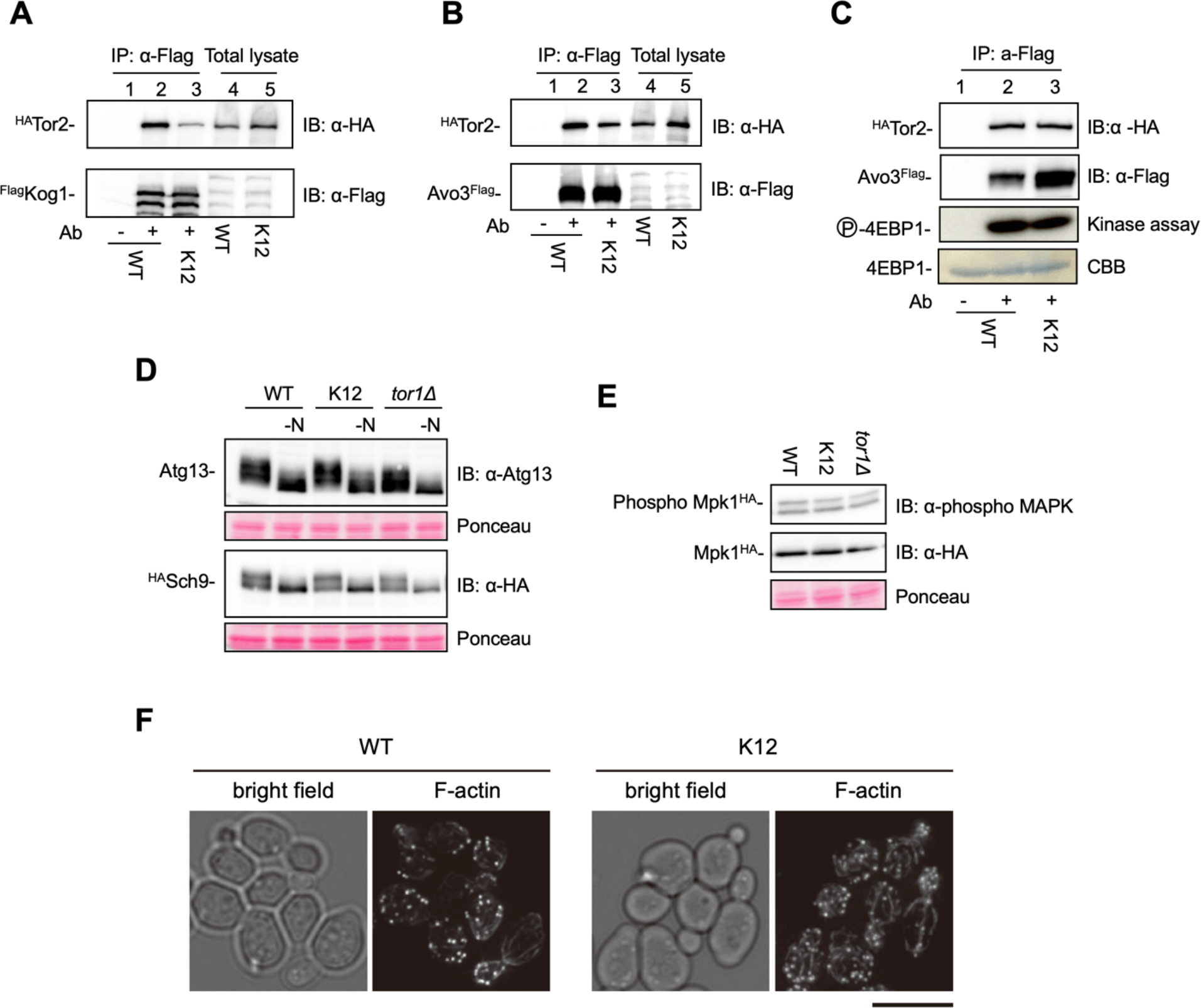
Tor2(K12) mutant almost loses TORC1 assembling ability and retains TORC2 assembling ability. **A.** Formation of Tor2-TORC1. ^Flag^Kog1 was immunoprecipitated from cell lysate of the wildtype and *tor2*(K12) (lanes 4 and 5), and co-precipitated ^HA^Tor2 in the immunocomplexes were detected (lane 1-3, lane 1 is immunoprecipitation control without antibody). *tor2*(K12) has almost no TORC1 assembling ability. **B.** Formation of Tor2-TORC2. ^Flag^Avo3 was immunoprecipitated from cell lysate (lanes 4 and 5), and co-precipitated ^HA^Tor2 contained in the immunocomplexes was detected (lanes 1-3). *tor2*(K12) retains the TORC2 forming ability. **C.** *In vitro* RI kinase assay of TORC2. The same amount of TORC2 was immunoprecipitated from cell lysate (estimated by the amount of ^HA^Tor2 protein), and TORC2 kinase assay was carried out using 4EBP1 as a substrate. TORC2 kinase activity of Tor2(K12) is comparable with that of the wild-type Tor2. Non-RI kinase assay also yielded similar results (Appendix Fig. S2). **D.** *In vivo* TORC1 kinase activities. Cells harbouring ATG13 (top) or ^HA^SCH9 (bottom) plasmid growing in YEPD at 30 °C were treated by nitrogen starvation (-N) for 30 min, and the phosphorylation state of Atg13 and Sch9 was examined by immunoblotting. TORC1 in cells has similar activity to that of the wildtype. **E.** *In vivo* TORC2 kinase activities. Cells harbouring MPK1^HA^ plasmid grown in YEPD at 30 °C were examined phosphorylation state of Mpk1 (top panel) by immunoblotting. TORC2 activity of Tor2(K12)-TORC2 in cells is also comparable to that of the wildtype. **F.** Localization of F-actin in the *tor2*(K12) mutant. WT and *tor2*(K12) mutant cells were grown in YEPD at 30°C and processed for F-actin staining. Bright-field images are shown on the left. Scale bar, 5 µm.

Next, we examined the cell phenotypes of the *tor2*(K12) and *tor1*Δ strains in the presence of TORC1 inhibitors (Fig. 5A). As previously reported, the *tor1*Δ strain is more sensitive than the wild-type strain to rapamycin, a selective inhibitor of TORC1, and caffeine, an inhibitor of TORC1 (Reinke *et al*, 2006; Sekiguchi *et al*, 2014), than those of the wildtype. Interestingly, the *tor2*(K12) strain had a different phenotype from the *tor1*Δ strain and was even more sensitive to rapamycin and caffeine than the *tor1*Δ strain. The sensitivity of the *tor1*Δ strain can be explained by a decrease in the total amount of TORC1. However, the hypersensitivity of the *tor2*(K12) strain cannot be explained by a decrease in TORC1, because the amount of Tor2-TORC1 is generally lower than that of Tor1-TORC1 (Loewith *et al*., 2002; Reinke *et al*., 2004). This result indicated that Tor2-TORC1 is distinct from Tor1-TORC1 in terms of its response to TORC1 inhibitors. Moreover, under several pH conditions, growth of the *tor2*(K12) and *tor1*Δ strains was observed (Fig. 5B). The *tor1*Δ cell grew better than the wildtype at even high pH (pH 8.0∼8.5). In contrast, *tor2*(K12) cells grew poorly at a high pH (pH 8.0) and did not grow at higher pH (pH 8.5). This result also indicates that Tor1-TORC1 has different role from Tor2-TORC1.

**Fig. 5.**
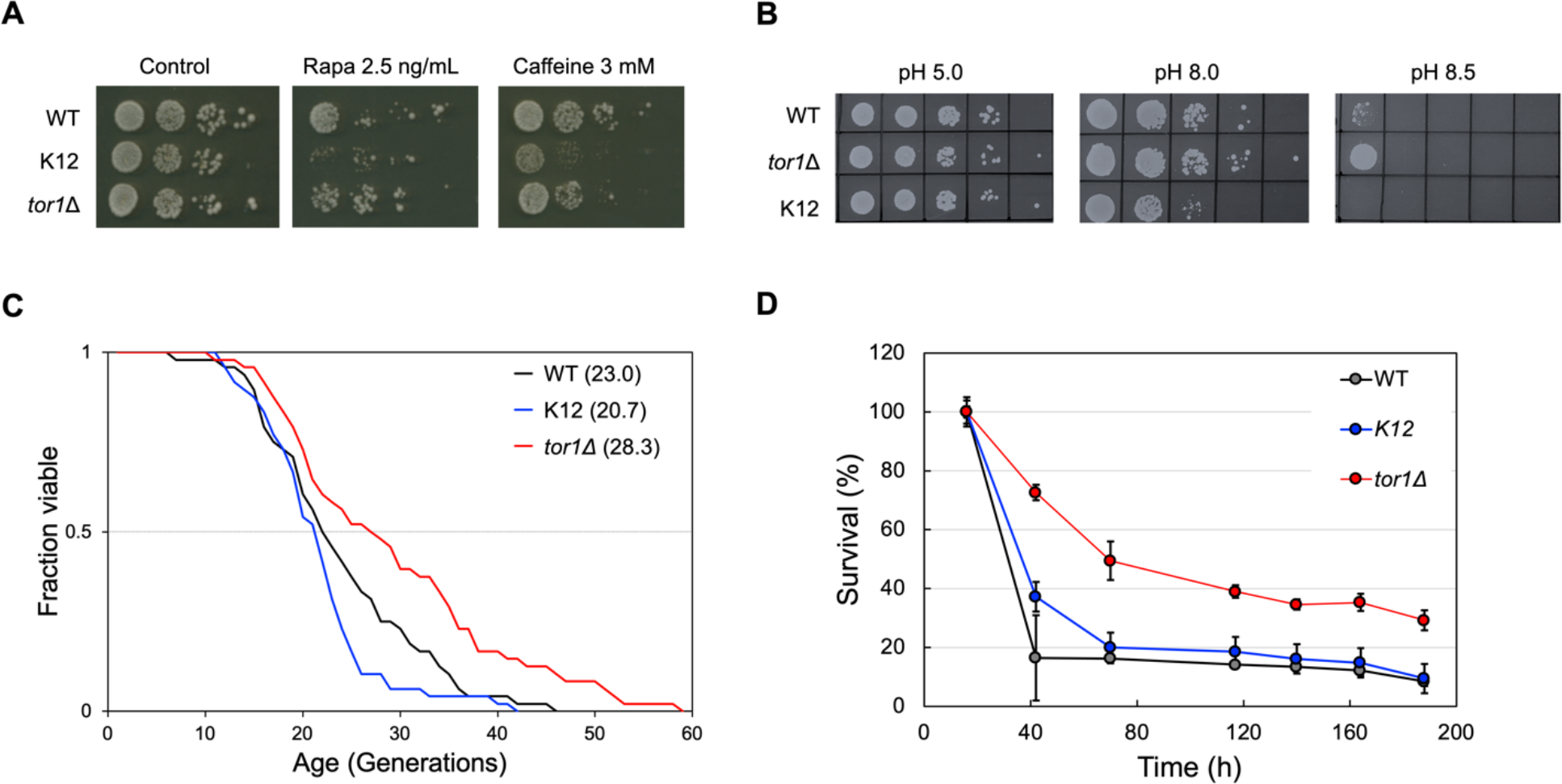
Cell phenotypes induced by designing Tor2, K12. **A.** Cell growth at various conditions, i.e., rapamycin and caffeine treatments. *tor2*(K12) strain has higher sensitivity for 2.5 ng/ml rapamycin (Rapa), a selective inhibitor of TORC1, and 3 mM caffeine, an inhibitor of TORC1. **B.** Cell growth at indicated pH conditions (pH 5.0, 8.0, and 8.5). *tor1*Δ strain is viable at higher pH conditions and K12 strain is only viable at lower pH conditions, compared to the wildtype. **C.** Replicative lifespans of the wild-type (Black), *tor2*(K12) (Blue), and *tor1*Δ (Red) strains. Mean life spans are shown in parentheses. Wilcoxon test, *p* = 0.030 (*tor1*Δ), *p* = 0.20 (K12); weighted log-rank test, *p* = 0.0054 (*tor1*Δ), *p* = 0.065 (K12) (versus wildtype). **D.** Chronological lifespans of the wild-type (Black), *tor2*(K12) (Blue) and *tor1*Δ (Red) strains. Quantitative data in the figures represent the average ± standard deviation (*n* = 3). Deletion of Tor1-TORC1 extends the lifespans of both, while the deletion of Tor2-TORC1 has a small effect on their lifespans.

Finally, replicative and chronological lifespans were measured for the wild-type, *tor1*Δ and *tor2*(K12) strains (Fig. 5C and D). The *tor1*Δ strain had a longer replicative lifespan than that of the wild-type strain, as previously reported (Kaeberlein *et al*., 2005). The replicative lifespan of the *tor2*(K12) strain was similar to that of the wildtype, although it seemed slightly shorter. The mean life spans of the wild-type, *tor1*Δ and *tor2*(K12) strains are 23.0, 28.3, and 20.7, respectively. All strains had chronological lifespans similar to their replicative lifespans: chronological lifespan of the *tor2*(K12) strain was similar to that of the wild-type strain, while the *tor1*Δ strain had longer chronological lifespans than the wild-type strain. Both lifespan results suggest that the roles of Tor1-TORC1 and Tor2-TORC1 are different from each other. It is possible that the *tor2*(K12) strain is less affected because of the lower amount of Tor2-TORC1. However, since the TORC1 activity itself is almost the same in both the same in both the *tor2*(K12) and *tor1Δ* strains (Fig. 4D), it is more likely that Tor1-TORC1 and Tor2-TORC1 contribute in different ways to lifespan regulation.

These phenotypic observations indicate that two types of TORC1s, namely Tor1-TORC1 and Tor2-TORC1, not only have common and essential functions, but also have distinct functions.

## Discussion

We engineered *S. cerevisiae* Tor2 based on computationally modelled Tor2-TORC1 and TORC2 structures. Through various cell biology and biochemical experiments, it was verified that the Tor2(K12) mutant maintains TORC2 activity but does not have TORC1 activity, as designed. Because only TORC1 activity is deleted from Tor2, the *tor2* mutant provides a strain without Tor2-TORC1 function, which is not created by the deletion of a gene because both TORC1 and TORC2 are essential for cells. By comparing the phenotypes of the *tor2* mutant strain with the *tor1*Δ strain, we found Tor2-TORC1 has distinct functions from that of Tor1-TORC1. Further research, for example, by solving and comparing high-resolution structures, could uncover the differences between Tor1-TORC1 and Tor2-TORC1 in detail.

In this study, we successfully altered the combinations of constituent proteins in a protein complexes using structure-based engineering, which has contributed to uncovering the biological function of the protein complexes. Various proteins form complex states in cells, many of which alter the combination of constituent proteins and exert their functions at the correct place and time. Our approach, which artificially engineered a combination of constituent proteins, enabled us to control celllar phenotypes and uncover the roles of protein complexes. However, a reasonable structural model is essential for the structure-based engineering. In this study, we designed mutants based on model complex structures predicted computationally by homology modeling and not on the experimental structures. Even if an experimental structure of the homologue protein is not available, AlphaFold 2 (Jumper *et al*., 2021), a high-accuracy protein structure prediction program using deep learning, has been developed, and we can use high-quality structural data for any proteins that we are interested in designing; note that experimental structures are better for designing mutants when they have been solved at the high resolution. Therefore, our approach applies to any target protein, and we could engineer native proteins based on their structure and uncover their biological functions.

The phenotypic difference between the K12 and *tor1*Δ strains not only indicates that the role of Tor1-TORC1 is not the same with that of Tor2-TORC1 but also provides more detailed information about the characteristics of Tor1-TORC1 and Tor2-TORC1. The higher sensitivity of the *tor2*(K12) mutant strain to TORC1 inhibitors may indicate that Tor2-TORC1 has a lower binding affinity or a more robust response to the inhibitors than Tor1-TORC1. The pH dependence of cell growth might be related to the activation of vacuolar-type ATPase (V-ATPase), an ATP-driven proton pump, by TORC1. Transport across the plasma membrane by V-ATPase controls the cytoplasmic pH condition and deletion of V-ATPase causes to deceleration of growth at high pH conditions (Kane, 2006; Nelson & Nelson, 1990), suggesting that high V-ATPase activity might enable growth at high pH conditions. The activity of mammalian V-ATPase is dependent on the activation of mTORC1; the inactivation of mTORC1 regulates the formation of a complete and active complex state (Ratto *et al*, 2022). Therefore, it is expected that *S. cerevisiae* TORC1 activity also affects the activation of V-ATPase, and its inactivation induces cell growth under alkali pH conditions. Our results were consistent with this expectation; the *tor1*Δ strain (permanent inactivation of Tor1-TORC1) grew under high pH conditions. In contrast, the K12 strain (almost permanently inactivated Tor2-TORC1) exhibited the opposite phenotype. Although these results should be investigated in more detail; future research on these two strains may contribute to our understanding of the mechanism of V-ATPase activation by TORC1.

The findings obtained from this study could provide clues to research the evolution of Tor2. The longer lifespan of the *tor1*Δ strain is the same phenotype with inhibition of mTORC1 or *S. pombe* TORC1, as previously reported (Harrison *et al*., 2009; Rodríguez-López *et al*., 2020). The pH dependence of growth of the *tor1*Δ strain (no Tor1-TORC1) is expected to be induced by the function, probably corresponding to mTORC1. Therefore, *S. cerevisiae* Tor1-TORC1 corresponds to TORC1s from other species, as mentioned in the Introduction. However, our results indicate that the role of Tor2-TORC1 is not the same with those of Tor1-TORC1 in *S. cerevisiae* and TORC1s in other species; it is a characteristic TORC1 of *S. cerevisiae*. This result also provides us information on the evolution of Tor. A simple hypothesis is *S. cerevisiae* Tor1-TORC1 is a special and later emergent one in the process of evolution because *S. cerevisiae* Tor2 forms both TORC1 and TORC2, similar to mTOR. However, our results indicate that Tor2-TORC1 is unique and a simple hypothesis of the evolution process is not reasonable. By further investigation of Tor2-TORC1, we could approach the evolution of Tor2.

We also obtained important findings regarding lifespan researches. Lifespan experiments showed that the *tor2*(K12) strain (almost no Tor2-TORC1) had a lifespan similar to that of the wildtype, although the *tor1*Δ strain (no Tor1-TORC1) had a longer lifespan. This result suggests that Tor1-TORC1 and Tor2-TORC1 had shortened and unaffected lifespans, respectively. By further investigating the differences between Tor1-TORC1 and Tor2-TORC1, we can uncover *S. cerevisiae* Tor2-TORC1’s function in more detail. This knowledge might enable us to create a mTORC1 similar to *S. cerevisiae* Tor2-TORC1 and to extend the lifespan by replacing the mTORC1 with a novel one in mammals without inhibiting the activity of mTORC1. This will enable the control of the lifespans of mammals.

## Materials and Methods

### Design protocol for Tor2 mutant

The model structures of Tor2-TORC1 and TORC2 were created using following procedure. Homology models of Tor2, Kog1, and Avo3 were individually generated using the SWISS-MODEL server (Waterhouse *et al*, 2018). The model structures were superimposed on the mTORC1 (PDB ID: 6BCX) and mTORC2 (PDB ID: 6ZWM) cryo-EM structures. To remove crashes between the atoms of each component, the components were shifted slightly from each other and, the side chains were repacked using the Rosetta protein design software (Leaver-Fay *et al*, 2011). These two model complex structures were compared and candidate residues for mutations that contribute to binding to Kog1 in Tor2-TORC1 and not to Avo3 in TORC2 were selected. The hydrophobic residues of the candidates were mutated to a larger hydrophilic but uncharged amino acid, glutamine. Positively or negatively charged residues were mutated to larger amino acids with the same charge, for examples, Lys to Arg. Several combinations of the candidate residues were experimentally validated.

### Strains, plasmids, media and genetic methods

The yeast strains, plasmids and DNA primers used in this study are listed in Appendix Tables S1, S2, and S3. Standard techniques have been used to manipulate yeast (Kaiser, 1994; Longtine *et al*, 1998). Antibodies against HA-epitope (16B12, COVANCE, x5,000 dilution), Flag-epitope (M2, Sigma, x5,000 dilution), phospho-p44/42 (#9101, Cell Signaling, x3,000 dilution), and thiophosphate ester-specific RabMb (Abcam, x5,000 dilution) were used as primary antibodies for immunoblotting at the indicated concentrations. Antibodies against Atg13 (x3,000 dilution) were used as described previously (Kamada *et al*, 2000). The FLAG-tagged AVO3 strain was generated following a previously described protocol (Longtine *et al*., 1998; Sung *et al*, 2005)

### Creating, cloning, and cell-based assay for *TOR2* mutants

*TOR2* mutants were generated and cloned using the designated gap repair cloning (GRC) method (Chino *et al*, 2010; Ma *et al*, 1987). The overall procedure is presented in Appendix Fig. S5. First, site-directed mutated TOR2 fragments were amplified by PCR using specific DNA primers (Appendix Fig. S5A, Appendix Table. S3). Next, these (2 or 3) DNA fragments were mixed and transformed into yeast cells with a linearised pRS314 vector (Appendix Fig. S5B). The plasmids created by GRC were rescued from yeast transformants.

The resultant TOR2 mutant plasmids were transformed into *TOR1 tor2*Δ (YYK1411) and *tor1* Δ*tor2*Δ (YYK1412) strains, harbouring pRS316[TOR2] plasmid. The transformants were streaked onto 5-FOA plates to select a *ura3* cells which loses the URA3-maker wild-type TOR2 plasmid and harboured only the mutated TOR2 plasmid. Growth on 5-FOA plates was used to evaluate the functions of TORC2 (*TOR1 tor2*Δ strain) and TORC1 TORC2 (*tor1*Δ *tor2*Δ strain). A YEPD plate was used as a growth control.

As for the integration of *tor2*(K12) allele into the *TOR2* locus, the pRS314[TOR2(K12)] plasmid created above was cloned into BYP9689 (pBSBleMX), and HindIII-HindIII (1.9kb, encoding FAT-FRB-kinase domain) region of the insert was deleted to make pBSBleMX[TOR2(K12) H3Δ]. The resulting plasmid was linearised by BamHI digestion and transformed into the BY4741 strain to generate the *tor2*::BleMX::*tor2*(K12) mutant. The phenotype of this strain was examined as shown in Appendix Fig. S6. This strain (YYK1551) was used for lifespan assay.

### Immunoprecipitation of TORC1 and TORC2

Immunoprecipitation of the Tor complexes was performed as previously described (Kamada, 2017). For the TORC1 experiment, YYK1467 and YYK1530 (^HA^TOR2 ^Flag^KOG1 strains) and for TORC2 experiment, YYK1464 and YYK1528 (^HA^TOR2 AVO3^Flag^ strains) were used. Yeast cells grown in YEPD at 30 °C overnight were collected and resuspended in Z-buffer (50 mM Tris-HCl pH7.5, 1 M sorbitol) containing 0.01 mg/OD_600_ cells zymolyase 100T (Nacalai Tesque). These were converted to spheroplasts with 30 min incubation at 30 °C. The spheroplasts were harvested, washed by Z-buffer once, and suspended in 10 μl/OD_600_ cells ice-cold IP-buffer (1xPBS, 2 mM MgCl_2_, 1 mM Na_3_VO_4_, 7.5 mM *p*-nitrophenyphosphate (*p*NPP), 10 mM β-mercaptoethanol, 1% Tween-20), containing protease inhibitors (40 μg/ml leupeptin, 80 μg/ml aprotinin, 20 μg/ml pepstatinA, 200 μg/ml 4-(2-Aminoethyl) benzenesulfonyl fluoride hydrochloride (AEBSF), and 1 mM PMSF). The cell suspension was gently mixed and incubated on ice for 5 min to break spheroplasts. The lysate was centrifuged 15,000*xg* at 4 °C for 10 min twice, and the clear lysate (700 μl) was incubated with 15 μl of Dynabeads protein G (Invitrogen) bound with or without (control) 1 μl of anti-Flag antibody (M2, Sigma) 4 °C for 2 h with gentle rotation. The beads were transferred into fresh microfuge tubes and washed thrice with 1xPBS or Tes-buffer.

### *In vitro* TORC2 kinase assay

TORC2 kinase activity was evaluated using RI and non-RI kinase assays as described previously (Allen *et al*, 2007; Kamada, 2017; Kamada *et al*., 2005). The resultant immunocomplex was washed once, suspended in 24 μl of Tes-buffer (25 mM Tes-KOH pH7.25, 100 mM KCl, 10 mM MgCl_2_) containing 2 μg of the substrate (^His6^4EBP1), and preincubated at 30 °C for 5 min. In the RI kinase assay, the reaction was initiated by the addition of 3 μl of 2 mM [γ-^32^P]ATP (222 TBq/mmol Perkin Elmer) to the mixture (final concentration, 0.2 mM, 0.2 MBq/reaction), and the reaction mixture (final volume 30μl) was further incubated at 30 °C for 10 min. The reaction was terminated by adding of 15 μl of 4x SDS-PAGE sample buffer and incubating at 65 °C for 5 min. The samples (20 μl) were subjected to SDS-PAGE (12.5%), and the phosphorylated proteins were analysed using autoradiography and a BAS5000 (Fuji Film). In the non-RI assay, the reaction was initiated by adding 6 μl of 1 mM ATPγS (Abcam) to the mixture (final concentration, 0.2 mM), and the reaction mixture (final volume, 30μl) was further incubated at 30 °C for 20 min. The reaction was terminated by adding 3 μl of 250 mM EDTA. The protein in the reaction mixture was alkylated with 1.7 μl of 50 mM *p*-nitrobenzyl mesylate (*p*NBM, 2.7 mM, Abcam) at room temperature for 80 min. The sample was added to 12 μl of 4x SDS-PAGE sample buffer and incubated at 95°C for 2 min, and a 20 μl aliquot was subjected to SDS-PAGE (12.5%). The phosphorylated substrate was analysed by immunoblotting using thiophosphate ester-specific RabMb (Abcam) according to the manufacturer’s protocol.

### *In vivo* kinase analyses

Immunoblotting was performed as previously described (Kamada, 2017). Yeast cells harbouring YEp352[ATG13], YCplac33[^HA^SCH9], or YEp352[MPK1^HA^] grown in YEPD medium at 30 °C. For the nitrogen starvation treatment, cells were collected, washed thrice with distilled water, transferred to synthetic dextrose (SD) (-N) medium (0.17% yeast nitrogen base without ammonium sulfate and amino acids, 2% glucose), and incubated for 30 min. Cells (10 OD_600_ units) were collected and fixed with 100 μl of ice-cold alkaline solution (0.2 N NaOH and 0.5% ²-mercaptoethanol). After 5 min of incubation on ice, 10 μl of 1.8 M NaOAc pH5.2 and 1 ml of ice-cold acetone were added to the sample and incubated at −20 °C to precipitate the proteins. The protein samples were precipitated with a microfuge for 5 min, air-dried, suspended in 100 μl of SDS-PAGE sample buffer, and incubated at 65 °C for 15 min. The samples were thoroughly dissolved by sonication and subjected to sodium dodecyl sulfate-polyacrylamide gel electrophoresis. For immunoblotting, peroxidase-conjugated goat anti-rabbit IgG (H+L) or sheep anti-mouse IgG (H+L) (Jackson ImmunoResearch) was used as a secondary antibody (x10,000 dilution). Immobilon Forte Western HRP (Merck) and Light-Capture II (ATTO) were used for signal detection.

### Actin staining

Staining for actin was performed and observed as described previously (Kamada *et al*., 2005; Nakano *et al*, 2011). YEPD-grown cells were fixed for 30 min by the direct addition of 37% formaldehyde stock to a final concentration of 5%. Fixed cells were collected, washed with 1xPBS thrice, and then bodipy-phallacidin (Molecular Probes) was added for 2 h at room temperature to stain F-actin as previously described (Kaiser, 1994). Fluorescence and bright-field images were captured using a personal DV microscope (Applied Precisions). Fluorescence images were acquired in 20 serial sections along the z-axis at intervals of 0.2 mm. All images were 3-dimensionally deconvolved and stacked using the quick projection algorithm in the SoftWoRx software (Applied Precisions).

### Replicative life-span assay

The replicative lifespans was measured for the wild-type (BY4741), *tor1*Δ (YYK332), and *tor2*(K12) (YYK1551) strains. Replicative lifespan was assayed as previously described (Nakajima *et al*, 2020). Yeast cells were thawed from the frozen stock and streaked onto YEPD agar plates. After 2 days, a single colony was spread onto a YEPD agar plate, and the cells were grown at 30 °C overnight. The next day, cells were transferred again to a fresh YEPD agar plates and grown overnight. Using a micromanipulator, 48 cells were arrayed on a plate and allowed to undergo one or two divisions. Virgin cells were then selected and subjected to lifespan analysis. Except during manipulation, the plates were sealed with Parafilm, incubated at 30 °C during the day and stored at 4 °C at night to avoid excessive budding. Daughter cells were removed by gentle agitation using a dissecting needle and scored every 2 h. For each of the 48 cell lines, buds from each mother cell were counted for at least 3 days until the division of living cells ceased. The mean replicative lifespan and the *p*-value were calculated using the Wilcoxon rank-sum test and weighted log-rank test relative to the wild-type strain, BY4741.

### Chronological life-span assay

The same cell strains as those used in the replicative lifespan assay were used. To measure cell survival, cells were grown in SD liquid medium, sampled during each growth phase, and plated on yeast extract agar plates after dilution (Ohtsuka *et al*, 2021a). After 4–7 days at 30 °C, using colony-forming units, the number of viable cells in 1-mL aliquots of culture was determined and divided based on the cell turbidity at each sampling time. Cell growth was then monitored according to the turbidity determined using a Bactomonitor (BACT-550) equipped with a 600 nm filter (Nissho Electric).

To measure the chronological lifespan, the wild-type (BY4741), *tor1*Δ (YYK332), and *tor2*(K12) (YYK1551) strains were cultured in SD medium with 240 mg/L leucine, 80 mg/L uracil, 80 mg/L histidine, and 80 mg/L methionine (*n* = 3).

## Acknowledgements

We thank Dr. T. Maeda for helpful suggestions, Dr. H. Moriya for his technical advice, Dr. J. Broach for his generous gift of the TOR2 plasmid, Y. Ito and S. Kawai for technical assistance, and the members of the Tokai Tor Conference (ToToCo) for general discussions. We thank the NIBB Center for Radioisotope Facilities and the NIBB Trans-Omics Facility for their technical support. We also thank the National BioResource Project Yeast for the plasmid distribution. This work was supported by the NINS program for cross-disciplinary study (Grant Number 01312108 to Y. K., Y. O., A. Y. and, T. K.) and by the grant of Joint Research by the National Institutes of Natural Sciences (NINS) (NINS program No, 01112205 to Y. K., Y. M., H. O., Y. O., A. Y. and, T. K.) and by the Japan Science and Technology Agency (JST) Precursory Research for Embryonic Science and Technology (PRESTO, Grant Number JPMJPR20E6 to T. K.) and by a Grant-in-Aid for Scientific Research (C) from the Ministry of Education, Culture, Sports, Science and Technology of Japan (to HO) (JP21K05363).

## Author Contributions

T. K. conceived and conceptualized this project in discussion with Y. K., Y. O., and A. Y. Y. K. and T. K. designed the research in discussion with Y. O. and A. Y. T. K. performed to generate computational structure models and designed Tor2 mutations. Y. K. and T. K. prepared plasmid constructs. Y. K. performed cell-based screening, co-immunoprecipitation analyses, kinase assays and phenotype investigations for the TORC1 inhibitors. T. K. investigated phenotypes at several pH conditions. Y. K., Y. O., and A. Y. performed actin staining. C.U. and Y. M. performed replicative life-span assay. H. O. performed chronological life-span assay. Y. K. Y. O. A. Y., and T. K. wrote the initial manuscript. All authors discussed the results and contributed to the final manuscript.

## Data Availability

The yeast strains and plasmids used in this study are available from the authors upon request.

